# Metagenomic characterization of a harmful algal bloom using nanopore sequencing

**DOI:** 10.1101/2020.11.13.381525

**Authors:** Peter W. Schafran, Victor Cai, Hsiao-Pei Yang, Fay-Wei Li

## Abstract

Water bodies around the world are increasingly threatened by harmful algal blooms (HABs) under current trends of rising water temperature and nutrient load. Metagenomic characterization of HABs can be combined with water quality and environmental data to better understand and predict the occurrence of toxic events. However, standard short-read sequencing typically yields highly fragmented metagenomes, preventing direct connection of genes to a single genome. Using Oxford Nanopore long-read sequencing, we were able to obtain high quality metagenome-assembled genomes, and show that dominant organisms in a HAB are readily identified, though different analyses disagreed on the identity of rare taxa. Genes from diverse functional categories were found not only in the most dominant genera, but also in several less common ones. Using simulated datasets, we show that the Flongle flowcell may provide an option for HAB monitoring with less data, at the expense of failing to detect rarer organisms and increasing fragmentation of the metagenome. Based on these results, we believe that Nanopore sequencing provides a fast, portable, and affordable method for studying HABs.

## INTRODUCTION

Harmful algal blooms (HABs) pose serious threats to not only the aquatic biomes but also human health. HABs occur when toxic or ecosystem-disrupting species of algae and cyanobacteria rapidly increase in abundance in water bodies, leading to contamination of drinking water supplies and closure of recreational waterways and fisheries. The presence of HABs is associated with both acute toxicity, as well as chronic health issues such as cancer (Etheridge 2010, Gorham et al. 2020). The frequency of HABs has increased with warming water temperatures over the last few decades and may increase with additional change in temperature, acidification, nutrient load, and oxygenation of fresh and marine aquatic environments (Glibert et al. 2005). However, strong species and strain specific interactions between the environment and aquatic microbial communities make prediction of HABs difficult (Griffith and Gobler 2020), necessitating ongoing monitoring of species composition and presence of toxins and taste-and-odor compounds, such as microcystins, geosmin, and 2-methylisoborneol (Dietrich 2006). A survey of reservoir managers found that the control of algal blooms was their highest concern (Schafran 2005). Considerable costs are incurred to control HABs and their downstream effects, for example approximately $70 million spent over 10 years in Waco, TX (Dunlap et al. 2015) and $80 million spent annually by the US Army Corps of Engineers (Schafran 2005). Other estimates place costs of HABs at approximately $100 million dollars annually in the US (Hoagland and Scatasta 2006).

Metabarcoding and metagenomic approaches to studying HABs are increasingly used (Anderson 1995, Otten et al. 2016, Hennon and Dyhrman 2020, Hatfield et al. 2020), but as generally applied have some limitations. Molecular probes and PCR primers are designed with specific targets, and can introduce quantitative bias or may fail to amplify unknown members of a microbial community (Krehenwinkel et al. 2017). The large (i.e. 23S/28S) and small (i.e. 16S/18S) ribosomal subunits and ribosomal internal transcribed spacers (ITS) have been most commonly employed as a metabarcode (Lepère et al. 2000, Sherwood and Presting 2007, Stern et al. 2012, Gamez 2018, Zhang et al. 2020), though primers targeting toxin and taste-and-odor compound-producing genes have also been developed (Tsao et al. 2014, Suurnäkki et al. 2015, Legrand et al. 2016). Metabarcoding and PCR-targeted approaches can only provide taxonomic identity and presence/absence data for subjects of interest. Total metagenomic assembly allows discovery of additional taxa and genes, though the typical 150-500 bp Illumina reads limit the ability of assembly algorithms to link broad genomic regions. Library preparation and sequencing typically rely on instrumentation, such as thermocyclers, qPCR machines, and DNA sequencers, that are too large or unstable to function under field conditions. The time between sample collection and data analysis is often longer than the 24-36 hours that can be necessary to take action to prevent toxins and taste-and-odor compounds from infiltrating drinking water systems.

The Oxford Nanopore MinION device offers a unique solution to overcome the above limitations on HAB microbiome research. MinION is a third generation sequencing platform that can produce an average of 15 Gbp of data consisting of long (> 10 kbp) reads, is portable, and is powered by USB (Oxford Nanopore 2020). Compared to Illumina short-read sequencing platforms with estimated error rates from 0.1-0.5% (Pfeiffer et al. 2018), the Nanopore error rate is much higher at 5-15% (Rang et al. 2018), though this can be reduced to about 1% by polishing the assembled contigs with read sequences (Vaser et al. 2017). The MinION’s portability has made it a popular choice for DNA sequencing where controlled laboratory conditions are unavailable and/or real-time data are necessary, such as on Arctic/Antarctic expeditions (Johnson et al. 2017, Gowers et al. 2019), on the International Space Station (Castro-Wallace et al. 2017, Burton et al. 2020), in tropical rainforests (Menegon et al. 2017, Pomerantz et al. 2018), and in clinical settings (Wongsurawat et al. 2019). Oxford Nanopore recently released the inexpensive Flongle flow cell which allows for sequencing of up to 2 Gbp at a cost of <20% that of the MinION flow cell (https://nanoporetech.com/products/ accessed 10 August 2020). This may be valuable in applications where increasing the number of sampling events (e.g. long-term monitoring) is more valuable than deep sequencing depth.

In this study, we applied MinION to generate the first HAB metagenome on a water sample from Cayuga Lake, New York. To assess consistency of taxonomic annotations among the current tools on long-read DNA sequences, we evaluated out-of-the-box performance of Centrifuge (Kim et al. 2016), Kaiju (Menzel et al. 2016), Kraken2 (Wood et al. 2019), and BLAST+ (Camacho et al. 2009). We also created *in silico* datasets to mimic the Flongle flow cell output and evaluate its consistency with MinION data.

## MATERIALS AND METHODS

### Sampling

One surface water sample was collected from an ongoing HAB in Cayuga Lake at Taughannock Falls State Park in Tompkins Co., NY (approximately 42.5470 N, 76.5984 W) on July 15 2019. The water sample was split into 6 tubes (6ml each), and centrifuged at 21,000*g* for 10min to pellet bacteria. For each tube, DNA was extracted using E.Z.N.A. Plant DNA kit (Omega Bio-tek) following the manufacturer’s protocol. The DNA was then pooled and concentrated by AMPure XP beads to a total of 50μl for library prep.

### Sequencing and assembly

The Nanopore sequencing library was prepared using the Ligation Sequencing kit (SQK-LSK109) and sequenced on MinION R9 flowcell (FLO-MIN106D) for 60 hours. Signal files were basecalled with Guppy v3.1.5 (Oxford Nanopore) with the high accuracy flip-flop mode.

Reads were adapter trimmed with Porechop (https://github.com/rrwick/Porechop) and assembled using Flye v2.7 (Kolmorogov et al. 2019) in its metagenomic mode (--meta flag) with an estimated genome size of 50 Mb (based on a previous assembly of the data). Assembled contigs were polished with five iterations of Racon v1.4.3 (Vaser et al. 2017) followed by one round of polishing with Medaka (https://nanoporetech.github.io/medaka). Assembly graph structure was visualized with Bandage (Wick et al. 2015).

### Taxonomic assignment and gene annotation

Polished contigs were taxonomically annotated using four tools in an out-of-the-box fashion (i.e. without creation of custom search databases or extensive search parameter optimization) in order to most closely replicate usage by end users. BLAST+ was used with the NCBI *nt* database downloaded on 10 June 2020. Centrifuge was used with the NCBI nucleotide non-redundant database last updated 3 March 2018 available from its website (https://ccb.jhu.edu/software/centrifuge/). Kaiju was run with its *nr_euk* database, which is the most inclusive and includes a subset of the NCBI *nr* database with sequences belonging to archaea, bacteria, viruses, fungi, and microbial eukaryotes (https://github.com/bioinformatics-centre/kaiju). Kraken2 was used with its option to create the NCBI *nt* database (kraken2-build – download-library nt --db nt) which was constructed on 26 February 2020. Results were interpreted based on either the best annotation for each contig selected by each program (Centrifuge, Kaiju, Kraken2) or the result with the highest bitscore (BLAST+). NCBI taxonomy database files (nodes.dmp and names.dmp) were used to parse taxonomic relationships of annotations. Gene prediction was performed using MetaGeneMark (Besemer and Borodovsky 1999, Zhu et al. 2010), while ribosomal RNA genes were identified with barrnap 0.9 (https://github.com/tseemann/barrnap) and compared to SILVA 138 LSU/SSU Ref NR 99 databases (Quast et al. 2013, https://www.arb-silva.de/documentation/release-138/). rRNA sequences were aligned with MAFFT v7.464 (Katoh and Standley 2013) and phylogenies inferred with IQ-TREE v1.6.12 (Nguyen et al. 2015). Genes were annotated into clustered orthologous groups (COGs) with eggNOG v5.0 (Huerta-Cepas et al. 2019). BUSCO 4.3.1 was used to estimate completeness of gene annotation (Seppey et al. 2019).

### Flongle data simulation

To mimic output by the Nanopore Flongle flowcell, data were subset to approximately 2 Gbp by pseudo-randomly selecting reads using shuf (GNU coreutils 8.21). Ten subsets were generated and checked to ensure a similar length distribution to the complete dataset. Each subset was assembled, polished, and annotated with BLASTN as described above.

### Data Availability

Supplemental files available through Figshare (https://figshare.com/projects/Nanopore_HAB_metagenome/92567) include the polished metagenome (File S1, Predicted genes and their locations within the metagenome (File S2), amino acid sequences (File S3), and taxonomic annotations for each contig at each taxonomic level (File S4).

## RESULTS

Following adapter trimming, 3.74 million reads (9.35 Gbp) were retained with an N50 length of 3651 bp and a mean length of 2498 bp. Maximum read length was 110 kbp. Assembly and contig polishing produced a metagenome size of 48.9 Mbp consisting of 2183 contigs with an N50 of 90 kbp (Table 1; File S1).

**Table 1.** Summary statistics of assemblies and predicted genes. Flongle results represent mean (standard deviation) of all 10 subsets.

| **Assembly** | **Total Length (Mbp)** | **Number** | **Mean Length (bp)** | **Longest (Mbp)** | **Shortest (bp)** | **N50** | **N70** | **N90** |
| --- | --- | --- | --- | --- | --- | --- | --- | --- |
| MinION | 48.9 | 2183 | 22408 | 2.30 | 80 | 90015 | 42028 | 12372 |
| Flongle | 17.1<br>(0.441) | 965<br>(46.4) | 17773 (754) | 1.31<br>(0.625) | 67 (28) | 64864<br>(4341) | 37296<br>(1432) | 10570<br>(1369) |
| **Predicted Genes** | **Total Length (Mbp)** | **Number** | **Mean Length (bp)** | **Longest (bp)** | **Shortest (bp)** | **N50** | **N70** | **N90** |
| MinION | 12.3 | 66490 | 184.8 | 4721 | 18 | 252 | 166 | 90 |
| Flongle | 4.28<br>(0.103) | 23268<br>(720) | 184.2 (1.92) | 3379<br>(572) | 18 (0) | 255.3<br>(2.75) | 164.1<br>(2.02) | 89.1<br>(1.10) |

### Taxonomic Annotation

Taxonomic annotation varied considerably among the classification tools. The number of contigs that could not be classified or could only be classified at the root of the taxonomy ranged from 255 (Centrifuge) to 500 (Kaiju). Because the unclassified contigs tended to be short, these represent only 1.5% (Centrifuge, Kraken) to 15% (Kaiju) of the total length of the assembly (Figure S1). At the superkingdom level, the majority of classified contigs were identified as Bacteria (range 81-90%), while Eukaryota were represented by 10-18% of contigs. Archaea and viruses made up a very small proportion of the results, representing less than 0.5% of classified contigs. At the genus level, the number of taxa identified ranged from 263 (Kaiju) to 309 (Centrifuge). Only 53 genera were shared among all classification tools, representing between 43% to 77% of the total assembly, and of these only 37 genera individually represent >0.5% of the contigs (Figure 1). In each case, over 50% of contigs were identified as *Anabaena* or *Dolichospermum* (here combined due to recent taxonomic revisions between these genera; Wacklin et al. 2009). Taxa shared similar proportions among tools at other taxonomic levels (Figure S2, File S4). For individual contigs, 27% were identified to the same genus by all tools, while for an additional 17% of contigs three of the tools agreed on one genus. For 20% of contigs, there was no agreement between any classification tools (including failure to classify at genus level). These ambiguously annotated contigs on average tended to be longer than the average length of other contigs, with an N50 length of 152 kbp vs 90 kbp, respectively. The number of contigs vs. total assembly size per genus showed a positive linear relationship (p < 2e^-16^, Kendall’s *τ* = 0.54).

**Figure 1.**
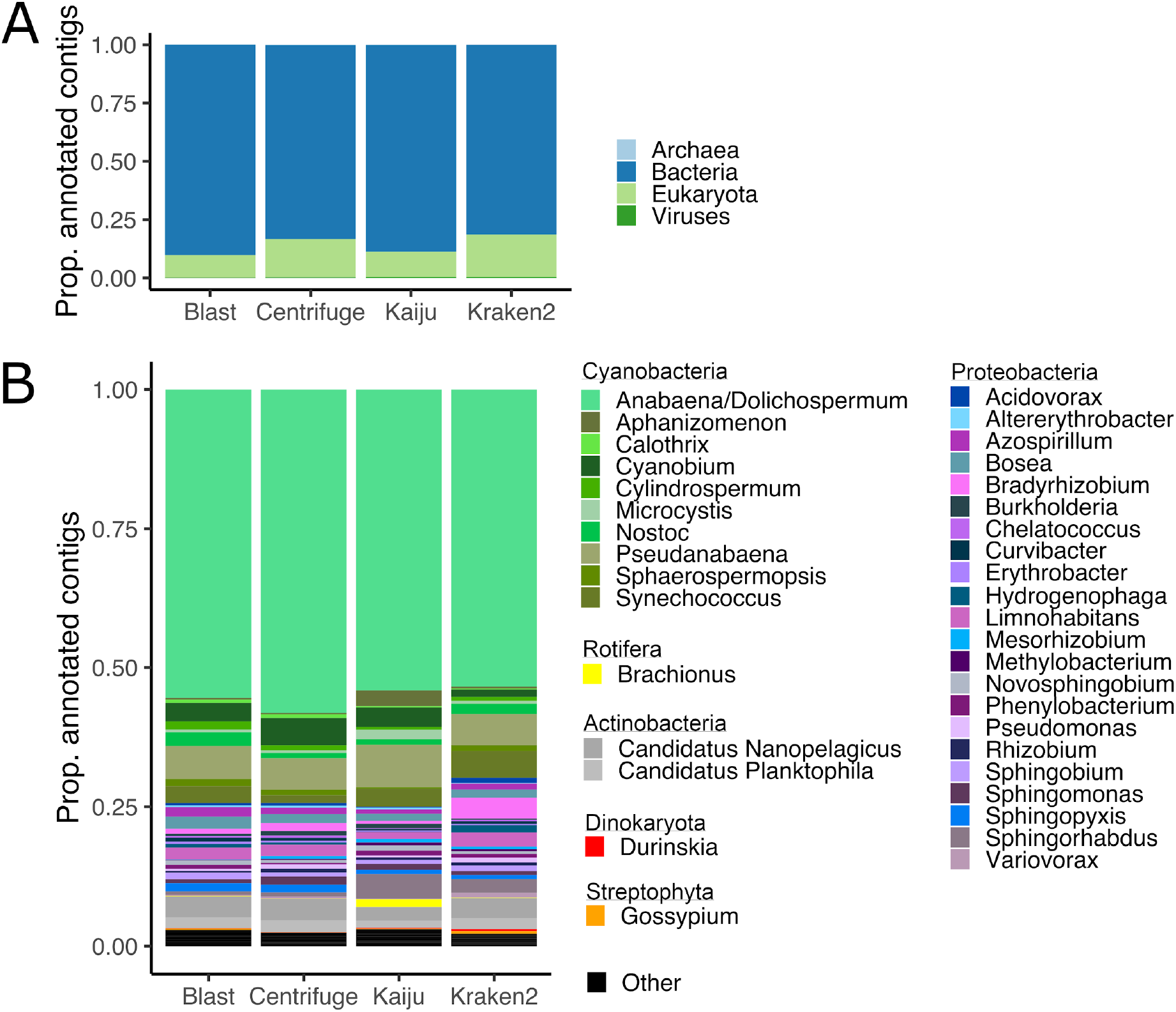
Proportion of contigs classified to superkingdoms (A) and genera (B) by each classification tool. Only genera identified by all classification tools that represent more than 0.5% of assembly shown. Individual genera present at <0.5% were combined into ‘Other’. Genera grouped by phylum.

Of the largest, high coverage cluster within the assembly graph—presumably representing *Anabaena/Dolichospermum* genome(s)—526 contigs form a relatively ‘knotty’ structure with many edges generally converging on two nodes (Figure 2). About 72% of contigs in this cluster were annotated as *Anabaena* or *Dolichospermum*. Coloration of edges by coverage suggests two levels, supported by Hartigan’s dip test of unimodality (p < 2.2e^-16^). This cluster may represent two genomes from closely related species, but attempts to isolate and reassemble them into more contiguous genomes were unsuccessful. There are 47 long contigs (>100 kbp) that are disconnected to the *Anabaena/Dolichospermum* cluster, which tend to have a lower coverage, except one 648 kbp contig annotated as *Anabaena/Dolichospermum* above 50X coverage. Three circular contigs with coverage from 390-628 X and lengths from 86-95 kbp were also annotated as *Anabaena/Dolichospermum*, and might be their plasmids.

**Figure 2.**
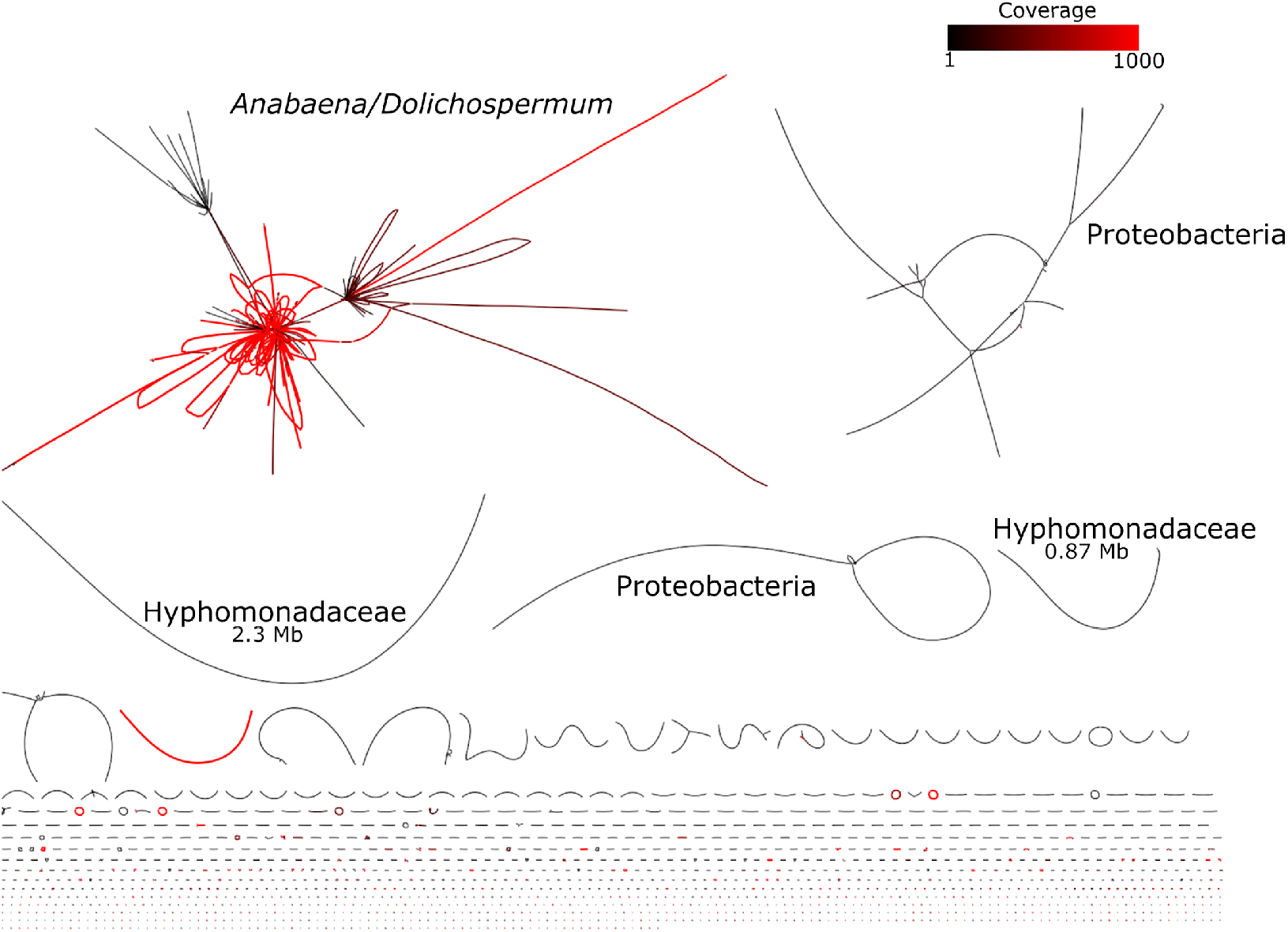
Assembly graph colored by coverage. Five largest clusters annotated with taxon at lowest level that provides majority agreement among all contigs. Contig lengths to scale.

### Gene Annotation

Gene annotation identified 66,490 genes with an N50 length of 756 bp (Files S2, S3). BUSCO analysis with the Bacteria v10 dataset found 99% complete genes in our metagenome (including 88% in the *Anabaena/Dolichospermum* cluster), suggesting this annotation is fairly robust. Genes were found representing every COG category (Figure 3). After genes with unknown function, the largest COG represented genes involved with amino acid transport and metabolism. Several COGs had very few representative genes, including: 1) RNA processing and modification; 2) chromatin structure and dynamics; 3) extracellular structures; 4) nuclear structure; 5) cytoskeleton. While the most abundant genera in the assembly also had the highest number of genes, several genera that were relatively rare in the assembly showed a high number of genes distributed across the predominant COG categories (Figure 3).

**Figure 3.**
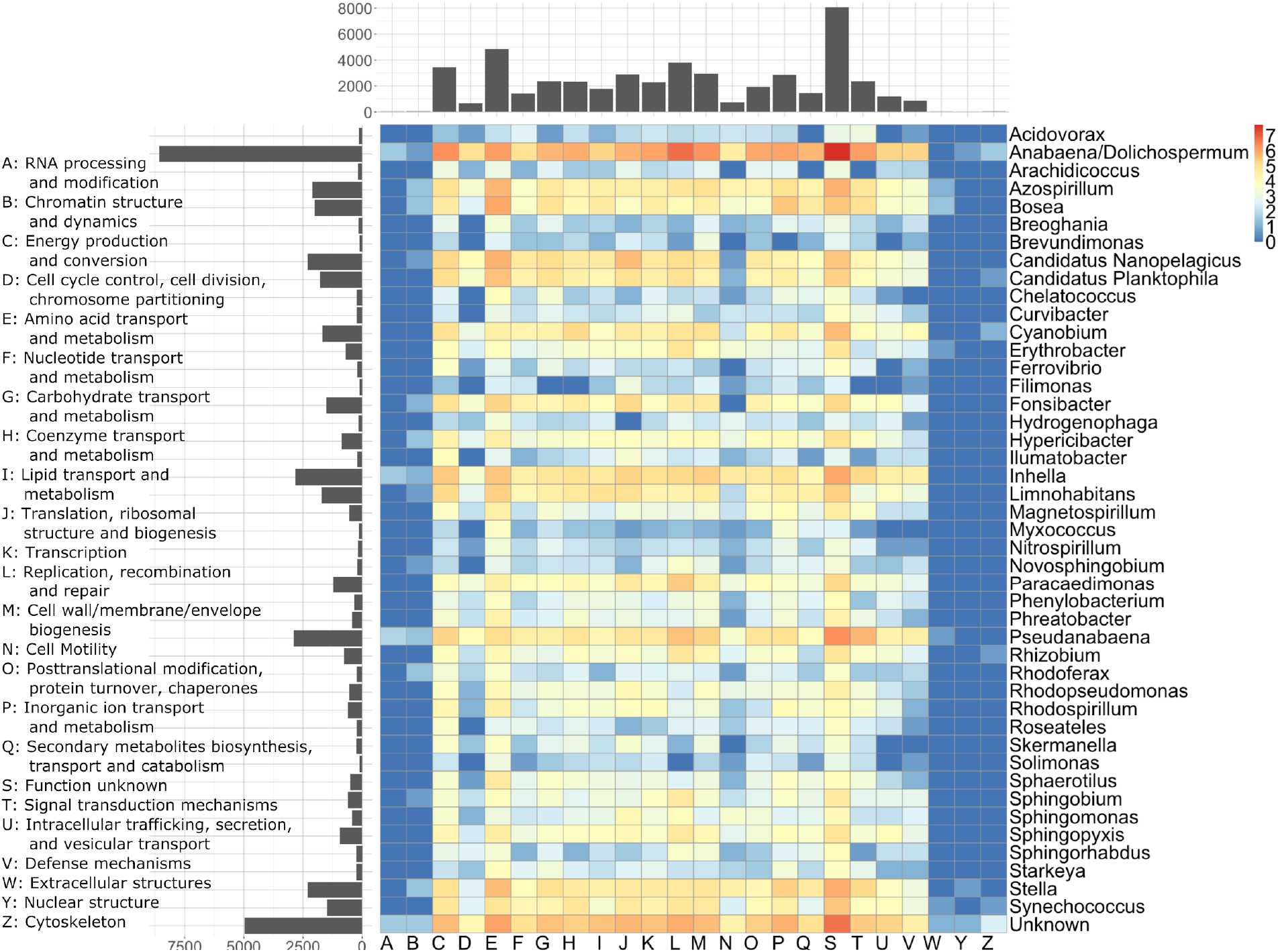
2D histogram of log-transformed COG annotations for genera with more than 100 annotated genes. Color legend represents log10-transformed number of genes. Histograms represent number of genes in respective row/column. X-axis categories described in left-hand legend.

Barrnap identified thirty-seven 16S and thirty-two 23S sequences with average lengths of 1185 bp and 2469 bp, respectively, though several fragmented sequences needed manual correction. Twenty-three 16S and fifteen 23S sequences matched entries in SILVA databases with greater than 95% identity and 95% coverage of the query sequence length. Phylogenies showed the most common rRNA sequences are nested in *Anabaena* (Figures S3, S4). Pairs of 16S and 23S from the same contig supported the presence of *Pseudanabaena, Candidatus* Nanopelagicus, *Candidatus* Fonsibacter, *Cercopagis pengoi*, and an unknown taxon sister to *Curvibacter, Rhodoferax*, and *Polaromonas*. In other cases 16S/23S pairs conflicted in their position to reference taxa. Several 16S/23S pairs nested within Eukaryota but with low similarity to all SILVA references may represent organellar rRNA sequences. However, these sequences also failed to match any sequence in NCBI *nt* with identity and alignment length above 90%.

### Flongle Data Simulations

The 10 data subsets created to mimic output of the Flongle flowcell resulted in assemblies with an average size of 17 Mbp (standard deviation = 0.4 Mbp) composed of an average of 965 contigs (standard deviation = 46). While each subset represented 21% of the total data, the subset assemblies averaged 35% of the size of the total assembly. Similarly, the number of genes and total size of the genes both averaged 35% as large as the total assembly.

Taxonomic annotation by BLAST was largely consistent at the superkingdom level, though a small percentage of the subset assemblies was identified as Archaea, which was not present in the total assembly (Figure 4). For other superkingdoms, the total assembly falls within the kernel density estimate of the subset results. At the genus level, larger discrepancies appeared (Figure 4), with similar trends apparent at other taxonomic levels (Figure S5). *Anabaena/Dolichospermum* was more highly represented in the subset assemblies than the total assembly, while most other genera showed the opposite trend. The range of subset results was large for some genera, such as *Sphingomonas* with a range of 15 percentage points, *Sphingopyxis* with 11 percentage points, and unannotated contigs with 11 percentage points.

**Figure 4.**
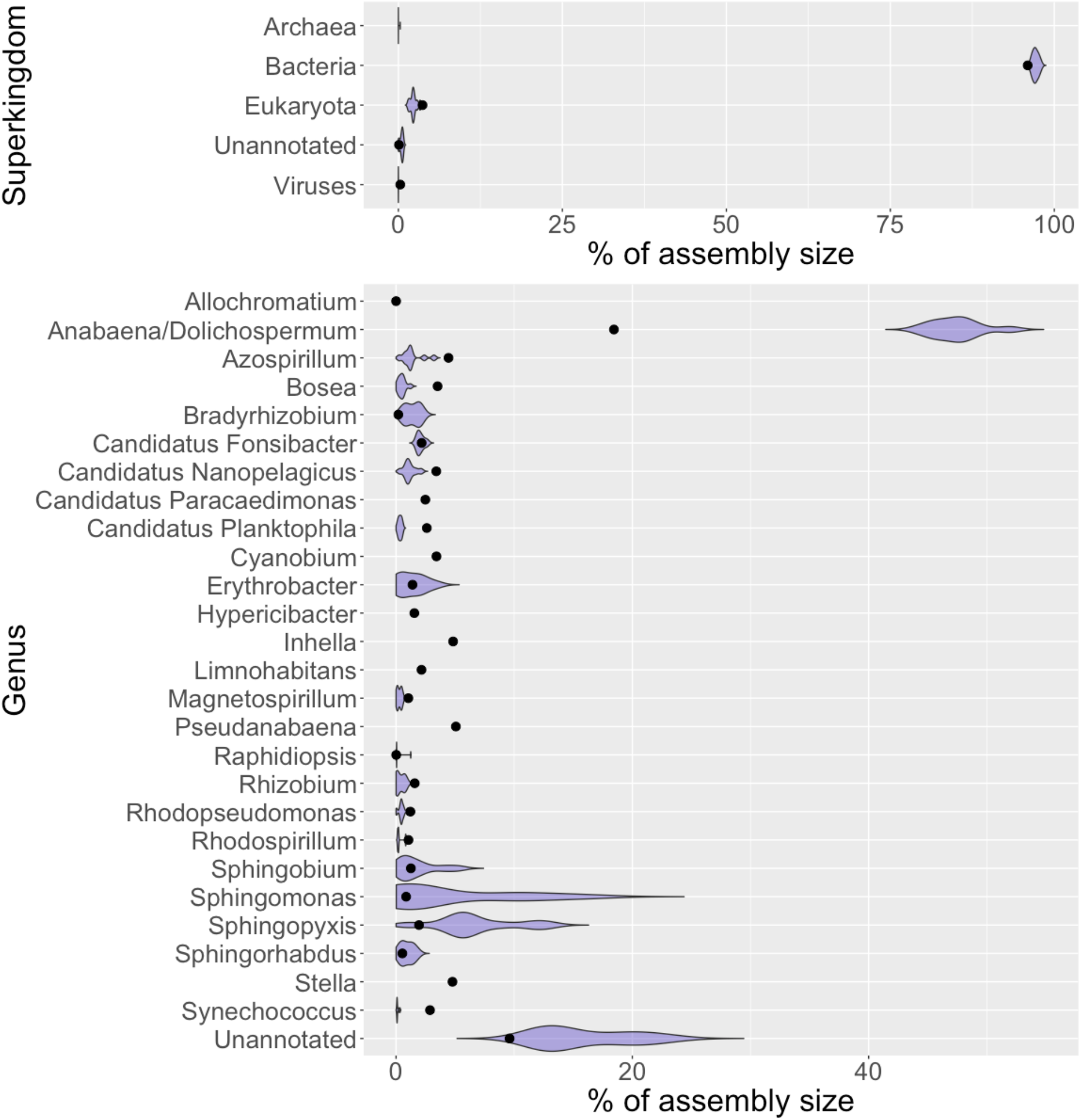
Comparison of assembly sizes for superkingdoms and genera between total assembly (dots) and Flongle-mimic subsets (violin plots) as annotated by BLAST. Only genera representing >1% of assembly shown.

## DISCUSSION

Our exploratory analysis using Nanopore DNA sequencing to characterize a harmful algal bloom highlights some potential uses for this technology, as well as some areas where caution should be used in interpreting the data. The main drawback of long-read DNA sequencing technology is a relatively high error rate compared to short-read sequencers. The error rate of raw sequences for the Nanopore flowcell and basecaller we used is estimated at roughly 10% (Wick et al. 2019), too high to be useful for k-mer based analysis or taxonomic annotation (data not shown). The development of Nanopore-focused assemblers that take into account the difference in coverage typical of metagenomic data, as well as polishing tools that remove errors through consensus calling and neural networks trained on specific models of Nanopore errors, provides methods to create highly accurate metagenomes without short-read sequences. These methods offer an advantage over short-read metagenomes by greatly increasing the contiguity of assembly with a small trade-off in accuracy. Our assembly returned many large contigs (N50 = 90 kbp), including one 2.3 Mbp contig identified as Hyphomonadacae bacterium that is about two-thirds the length of its most closely related bacterial genome in NCBI’s Genome database. We expected a more contiguous genome of the predominant cyanobacteria in our sample (*Anabaena* and/or *Dolichospermum*), but those contigs formed a few highly connected and structurally unresolved clusters. This can occur when a genome contains long repeat-rich regions or when multiple organisms share portions of their genomes that are similar. The structure of the assembly graph combined with significant bimodal distribution in coverage suggests that at least two highly similar organisms may be present in the *Anabaena/Dolichospermum* cluster, resulting in several shared edges connecting otherwise distinct clusters of contigs. However, our attempts to isolate and reassemble the raw data contributing to the *Anabaena/Dolichospermum* cluster did not improve the assembly.

BLAST, Centrifuge, Kaiju, and Kraken2 all were able to provide taxonomic annotations for a majority of contigs. Kaiju failed to annotate longer contigs than the others, possibly because it uses a smaller protein search database derived from NCBI *nr*, as opposed to the nucleotide-based *nt* used by the other tools. The low level of shared genus richness—where only 27% of contigs were placed in the same genus by every tool and roughly 20% of the genera identified by each tool are shared among all four—shows that caution needs to be exercised in determining community composition. While the more abundant organisms in this community may be identified with some certainty, contigs receiving mixed taxonomic annotations may need to be manually inspected. We processed BLAST results using the match between each contig and the reference database with the highest score to assign an identity to the contig. Individual matches tended to be short, with an average of 537 bp, so it may be more informative to examine all matches across different parts of the contig and use taxonomic distance (least common ancestor) between best hits to refine the overall identity, similar to how Centrifuge, Kaiju, and Kraken2 operate. Kraken2 provides an option to filter results by a user-defined confidence score, requiring a given fraction of a sequence to match a taxonomic level to receive an annotation. However, even at a low confidence level of 0.1, Kraken2 failed to provide many meaningful results from our data with 68% of contigs unclassified and of those classified, many were placed at the order level and above, suggesting that the application of this approach for metagenomics is limited.

Any classification is only as good as the reference database being searched. While NCBI *nt/nr* are the traditional databases for classification software, inclusion of the NCBI Whole Genome Shotgun (WGS) database can greatly improve results (Martí and Garay 2019). The short matches between our metagenome and *nt/nr* databases highlight the need to incorporate more genomic data for comparison against the increasing number of long-read metagenomic surveys. The software we tested all provide ways to create custom search databases, but this can require technical knowledge and resources many end users do not possess (e.g. our server with 768 GB RAM was memory-limited when trying to create a Centrifuge database from the current *nt* database). As a result, software that are not provided with regularly updated search databases, such as the most recent Centrifuge *nt* release from April 2018, quickly become obsolete as new genomic data are regularly added to the source databases. Additionally, updates in nomenclature, such as the transfer of some *Anabaena* into *Dolichospermum*, and persistence of polyphyly in bacterial phylogenies will confound attempts to use a least common ancestor approach to reconcile multiple matches (Wacklin et al. 2009, Cirés and Ballot 2016). This may partially explain why the proportion of contigs placed into *Anabaena* vs. *Dolichospermum* varies between BLAST/Kraken2 vs. Centrifuge/Kaiju, but in combination the genera are roughly equal. An independent sample from the same HAB event was microscopically identified to *Dolichospermum* (Bloom code 19-3435-B2, Community Science Institute, Ithaca, NY), supporting our decision to combine *Anabaena* and *Dolichospermum* results.

Our gene annotation and classification showed a broad diversity of genes obtained not only from the most abundant organisms, but also from many with lower abundance in our sample. This demonstrates one of the advantages of long-read sequencing in that much longer contigs allow connecting more genes to a single taxon, in addition to being able to taxonomically place previously unknown genes or genes without a known function. Efforts to incorporate genomic and proteomic data into HAB prediction models will be aided by this technologic development (Hennon and Dyhrman 2020). While we did not identify any in our sample, genes related to production of toxin and taste-and-odor compounds, such as geosmin synthase, 2-methylisoborneol (MIB) synthase, the anatoxin-a synthetase gene cluster, and the microcystin synthetase gene cluster, could be easily gleaned from this type of metagenome. This is consistent with failure to detect microcystin in this HAB at the drinking water threshold (<3 μg/L) by the Community Science Institute (Ithaca, NY). The ability of the Nanopore platform to also sequence RNA suggests that monitoring the expression of these genes could provide a real-time, field-based method to track the production of these dangerous compounds.

We created ten datasets *in silico* to estimate the potential of Oxford Nanopore’s Flongle flow cell to assemble and characterize a metagenome as well as the MinION flow cell that produced 5 times more data. The average Flongle assembly size was disproportionately large compared to the reduced size of the datasets, suggesting that some of the MinION data could be considered excess. However, the distribution of contig length shows the Flongle assemblies varied tremendously, with the length of the longest contig ranging from 2.5 to 0.6 Mbp and N50 ranging from 60 to 78 Kbp. Since the length distribution of the reads in each dataset was similar, this suggests the difference in assemblies is due to random assortment of reads derived from different organisms. While the proportion of the Flongle assemblies annotated to each taxonomic group was similar to the MinION assembly at the superkingdom level, increasing levels of taxonomic resolution showed the most prevalent organisms were overrepresented in the Flongle results, while the least prevalent organisms tended to be underrepresented. This is consistent with the methodology of the assembly and polishing software, where contigs with very low coverage are removed from the final assembly. Based on our results the Flongle may be useful for capturing large genomic fractions of the predominant members of an HAB, though it is more likely to miss minor taxa in the community.

### Conclusions

The profound effects of harmful algal blooms (HABs) on human health, economic activity, and ecological function compounded with the expectation that changes to the environment will alter the frequency and intensity of HABs necessitates a better understanding of the patterns and processes driving these events. Genomics provide complementary evidence to traditional data focused on taxonomic identification, quantification, and water quality measurements. Comparing four tools used for classifying DNA sequences, we found that assigning taxonomy with high confidence in our Nanopore-derived HAB metagenome was often difficult, though as more complete genomes are added to reference databases, we expect this to be resolved. We recovered a diversity of genes from many functional groups, even from organisms present at relatively low abundance in the metagenome. Our *in silico* datasets mimicking the Flongle flow cell for the Nanopore MinION and GridION sequencers suggests this may be a valuable tool for water quality managers who need to repeatedly monitor for HABs, with the tradeoffs being decreased detection of low abundance organisms and greater fragmentation of the metagenome. Given the portability, affordability, and short time from sample-to-sequence of the Nanopore platform, we believe this form of long-read sequencing shows potential for the study and monitoring of algal blooms.

## ACKNOWLEDGEMENTS

We thank Taughannock Falls State Park for access to the collection site and the Community Science Institute for sharing their data from this bloom event. V.C. thanks Duncan Hauser and Alaina Petlewski for laboratory training.

---

**Figure S1.**
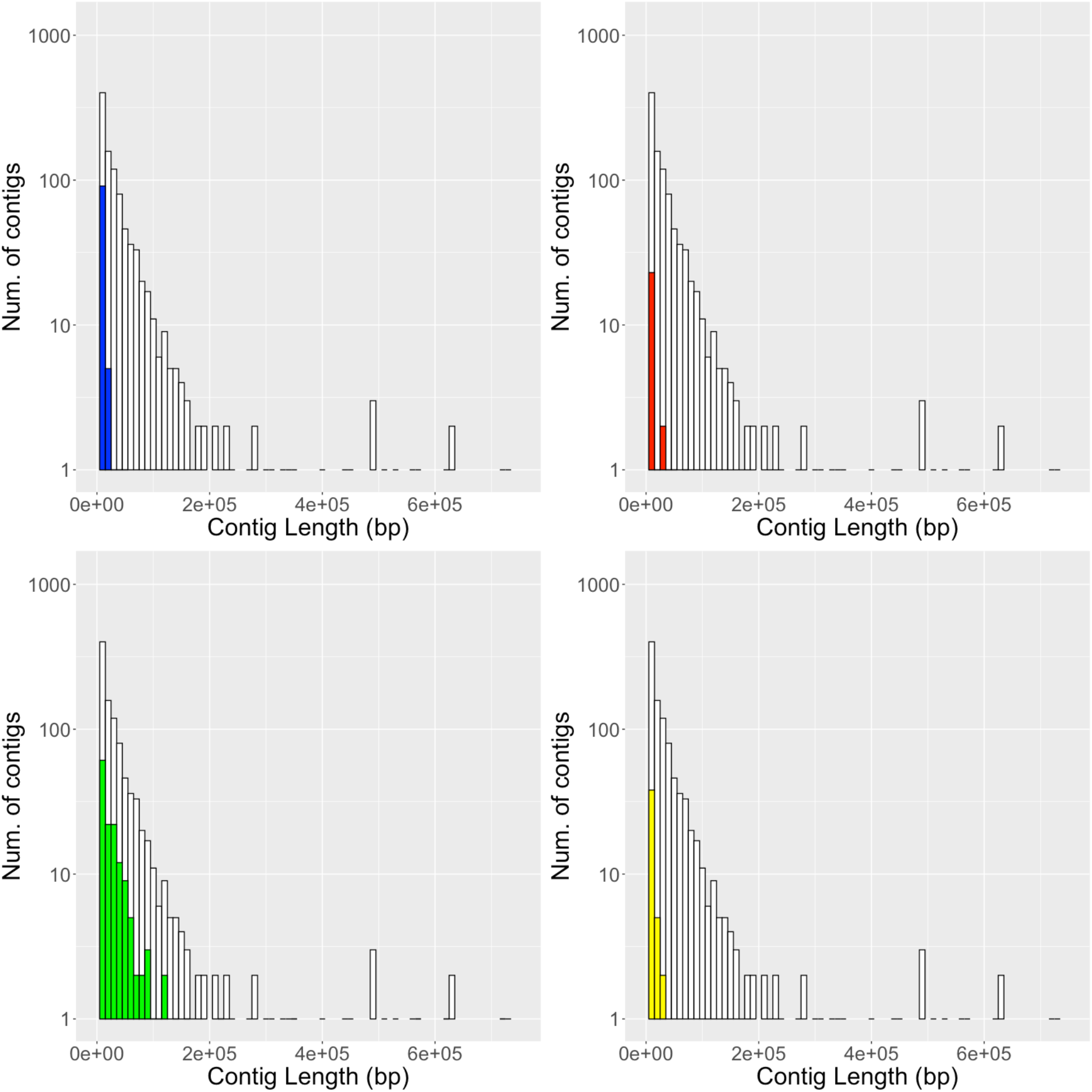
Histograms of unclassified contigs by length. Blue = Blast, Red = Centrifuge, Green = Kaiju, Yellow = Kraken2, White = all contigs.

**Figure S2.**
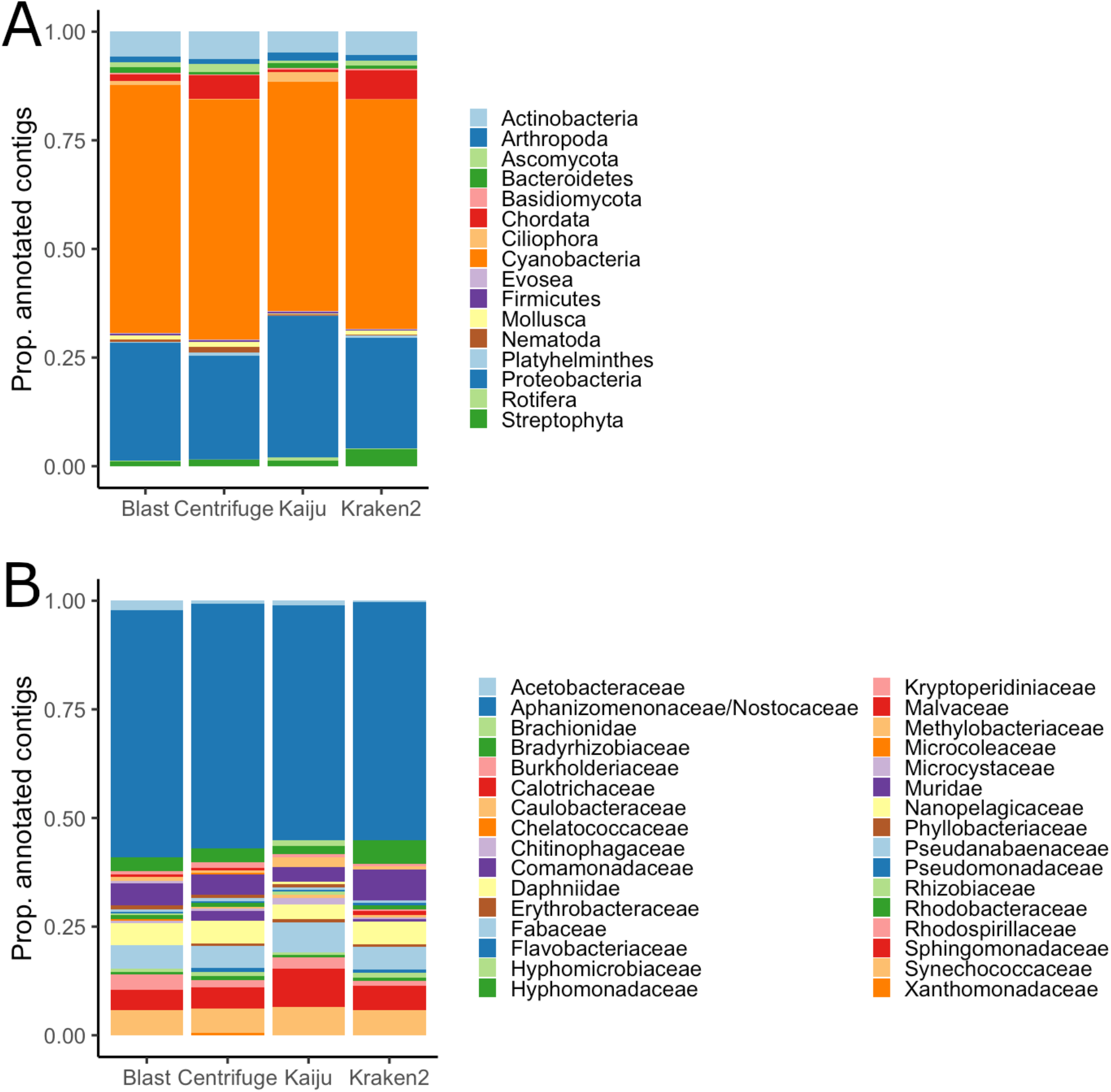
Proportion of contigs classified to phylum (A) and family (B) by each classification tool. Only taxonomic groups identified by all classification tools shown.

**Figure S3.**
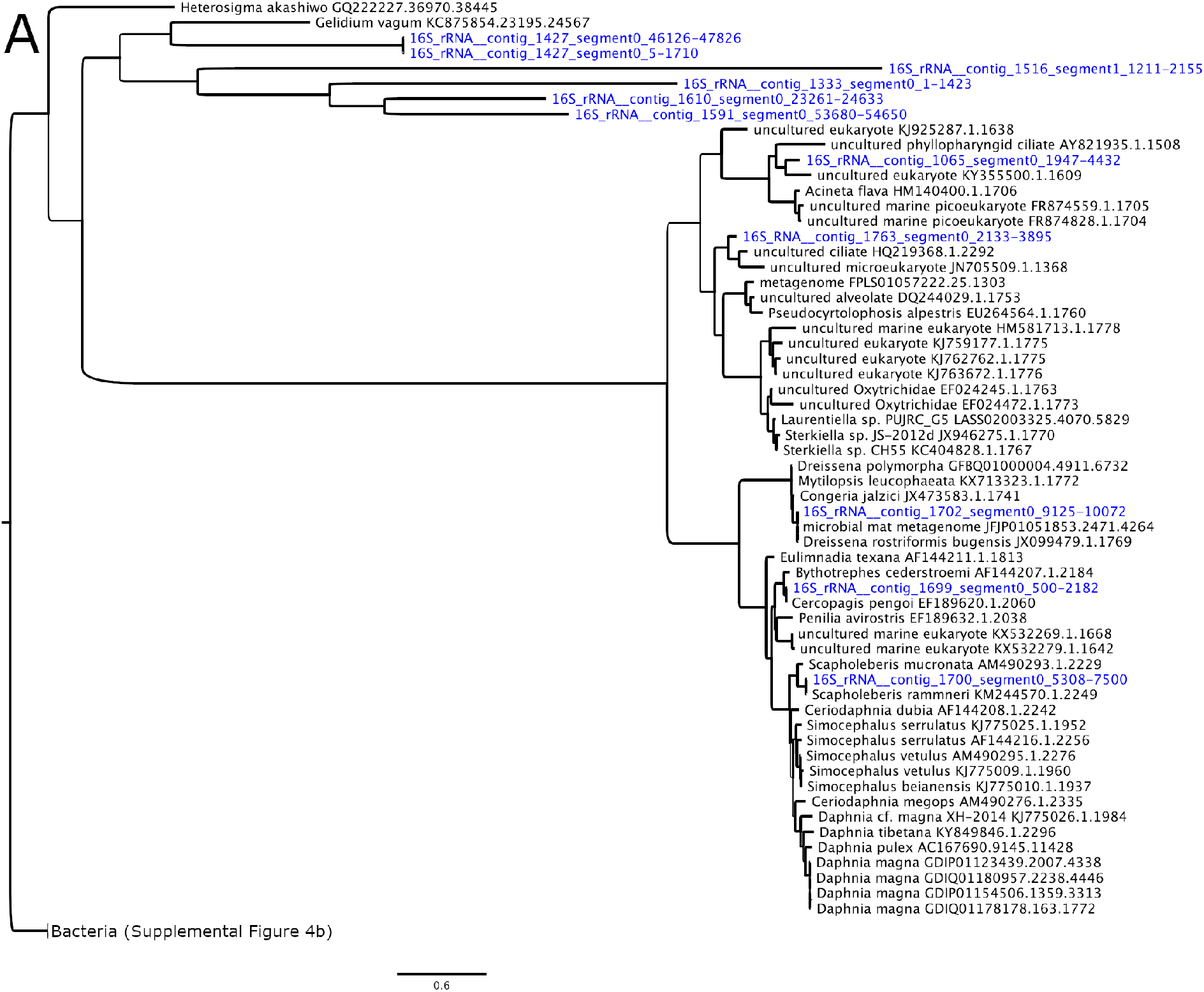
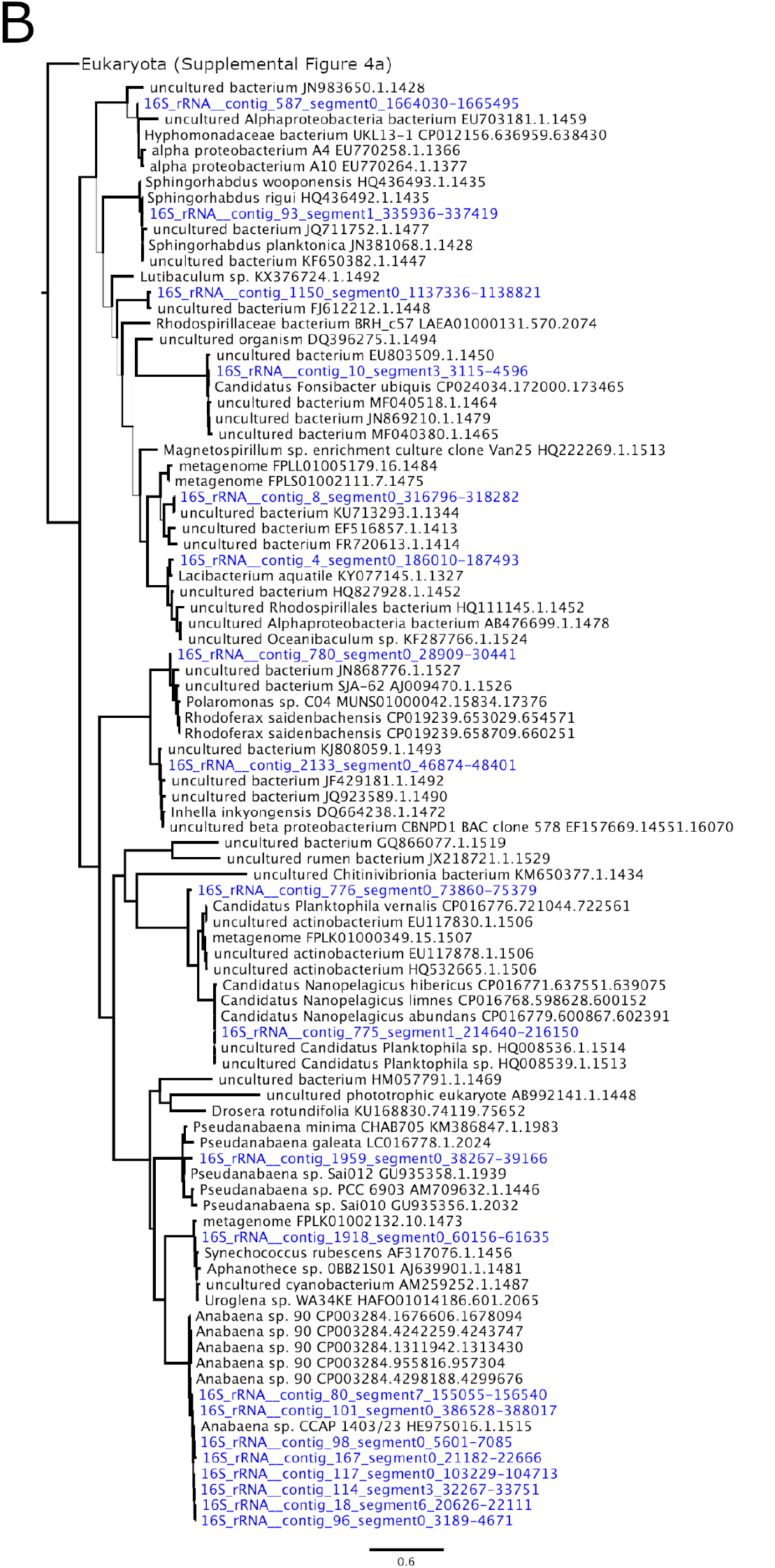
16S rRNA phylogeny of metagenome-recovered sequences (blue) and SILVA references (black). Tree rooted along the branch separating Eukaryota (A) and Bacteria (B). Branch thickness is proportional to bootstrap support with scale bar width = 100.

**Figure S4.**
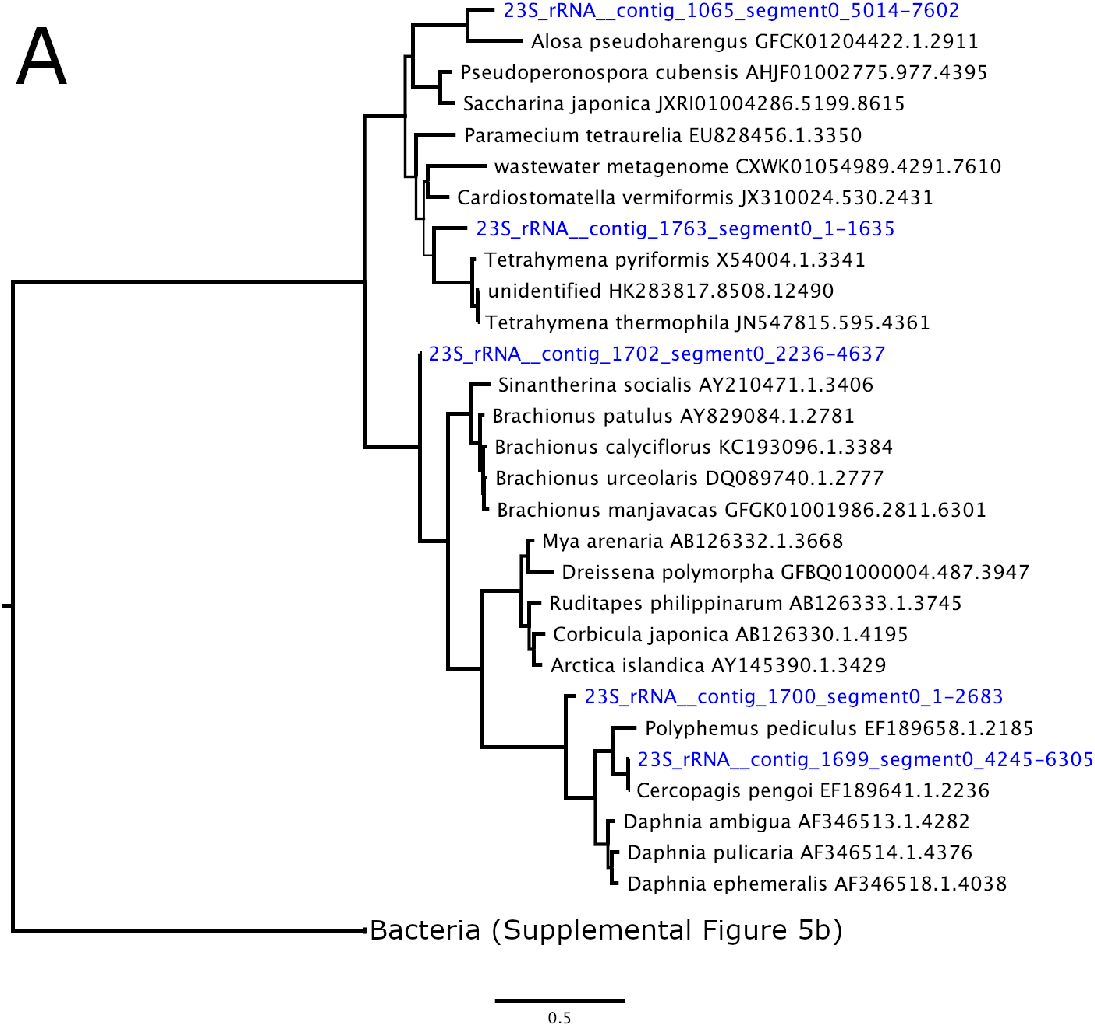
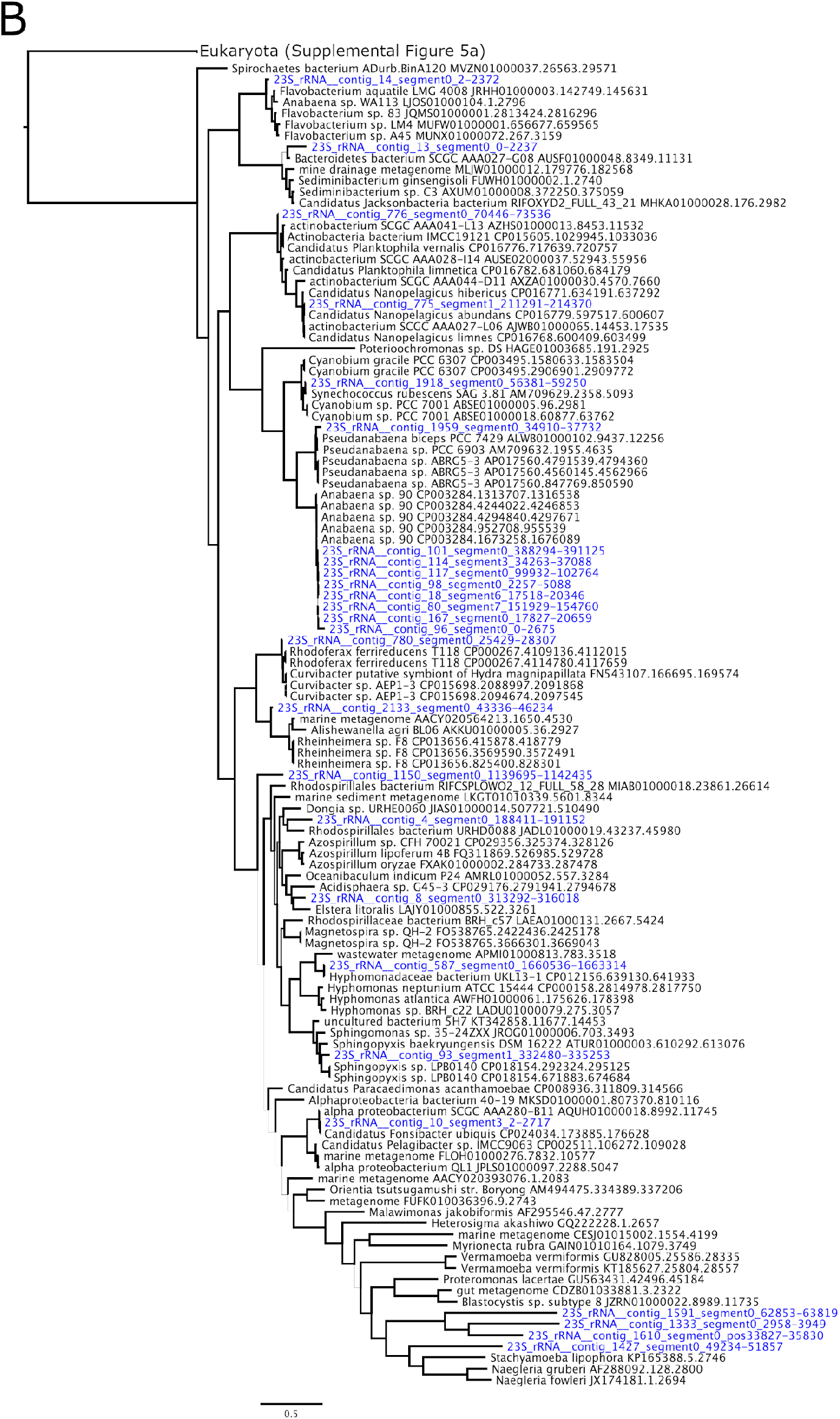
23S rRNA phylogeny of metagenome-recovered sequences (blue) and SILVA references (black). Tree rooted along the branch separating Eukaryota (A) and Bacteria (B). Branch thickness is proportional to bootstrap support with scale bar width = 100.

**Figure S5.**
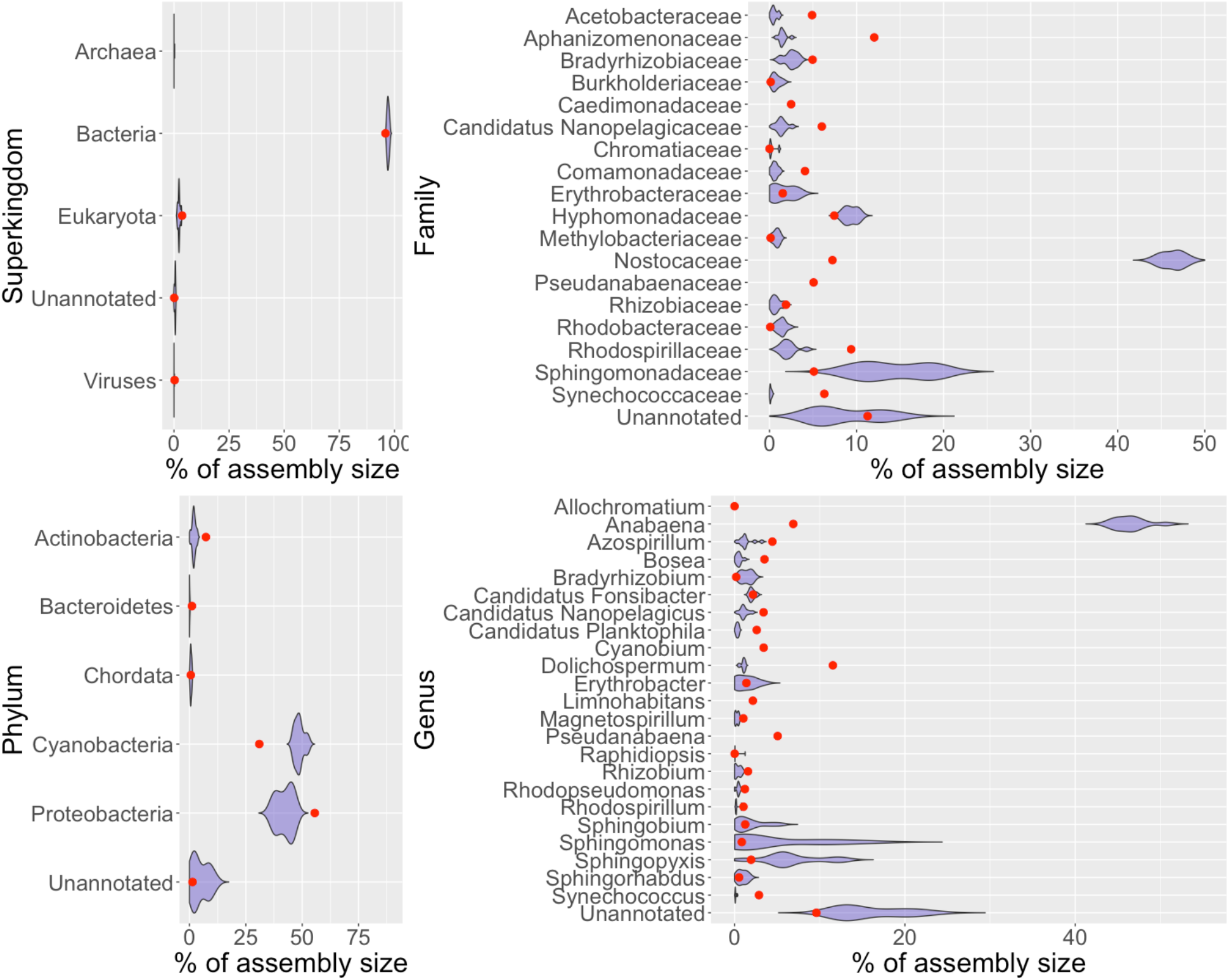
Comparison of assembly sizes for superkingdoms, phyla, families, and genera between total assembly (dots) and Flongle-mimic subsets (violin plots) as annotated by BLAST. Only taxonomic groups representing > 1% of assembly shown.

## REFERENCES

Besemer, J., and M. Borodovsky, 1999 Heuristic approach to deriving models for gene finding. Nucleic Acids Research 27(19): 3911–3920. https://doi.org/10.1093/nar/27.19.3911

Burton, A.S., S.E. Stahl, K.K John, M. Jain, S. Juul, et al., 2020 Off Earth identification of bacterial populations using 16S rDNA nanopore sequencing. Genes 11: 76.

Camacho, C., G. Coulouris, V. Avagyan, N. Ma, J. Papadopoulos, et al., 2009 BLAST+: architecture and applications. BMC Bioinformatics 10: 421.

Castro-Wallace, S.L., C.Y. Chiu, K.K. John, S.E. Stahl, K.H. Rubins et al., 2017 Nanopore DNA sequencing and genome assembly on the International Space Station. Scientific Reports 7: 18022. http://doi.org/10.1038/s41598-017-18364-0

Dietrich, A., 2006 Aesthetic issues for drinking water. Journal of Water and Health 4(S1): 11–16.

Dunlap, C.R., K.S. Sklenar, and L.J. Blake, 2015 A costly endeavor: addressing algae problems in a water supply. Journal of the American Water Works Association 107(5): E255–E262.

Etheridge, S.M., 2010 Paralytic shellfish poisoning: Seafood safety and human health perspectives. Toxicon 56(2): 108–122.

Gamez, T., 2018 The use of 23S ribotyping to detect harmful and nuisance phytoplankton in a large, subtropical reservoir during an extended drought period (Unpublished thesis). Texas State University, San Marcos, Texas.

Gilbert, P.M., D.M. Anderson, P. Gentien, E. Granéli, and K.G. Sellner, 2005 The global, complex phenomena of harmful algal blooms. Oceanography 18(2): 136–147.

Gorham, T., E.D. Root, Y. Jia, C.K. Shum, and J. Lee, 2020. Relationship between cyanobacterial bloom impacted drinking water sources and hepatocellular carcinoma incidence rates. Harmful Algae 95: 101801.

Gowers, G.-O. F., O. Vince, J.-H. Charles, I. Klarenberg, T. Ellis, et al., 2019 Entirely off-grid and solar-powered DNA sequencing of microbial communities during an ice cap traverse expedition. Genes 10: 902. https://doi.org/10.3390/genes10110902

Griffith, A.W., and C.J. Gobler, 2020 Harmful algal blooms: a climate change co-stressor in marine and freshwater ecosystems. Harmful Algae 91: 101590.

Hatfield, R.G., F.M. Batista, T.P. Bean, V.G. Fonseca, A. Santos, et al., 2020 The application of nanopore sequencing technology to the study of dinoflagellates: a proof of concept study for rapid sequence-based discrimination of potentially harmful algae. Frontiers in Microbiology 11: 844.

Hennon, G.M.M., and S.T. Dyhrman, 2020 Progress and promise of omics for predicting the impacts of climate change on harmful algal blooms. Harmful Algae 91: 101587.

Huerta-Cepas, J., D. Szklarczyk, D. Heller, A. Hernández-Plaza, S.K. Forslund, et al., 2019 eggNOG 5.0: a hierarchical, functionally and phylogenetically annotated orthology resource based on 5090 organisms and 2502 viruses. Nucleic Acids Research 47(D1): D309–D314. https://doi.org/10.1093/nar/gky1085

Hoagland, P., and S. Scatasta, 2006 The economic effects of harmful algal blooms, in Ecology of Harmful Algae. Ecological Studies (Analysis and Synthesis) vol 189, edited by E. Granéli and J.T. Turner. Springer, Berlin, Heidelberg. https://doi.org/10.1007/978-3-540-32210-8_30

Johnson, S.S., E. Zaikova, D.S. Goerlitz, Y. Bai, and S.W. Tighe, 2017 Real-time DNA sequencing in the Antarctic dry valleys using the Oxford Nanopore sequencer. Journal of Biomolecular Techniques 28(1): 2–7. https://doi.org/10.7171/jbt.17-2801-009.

Katoh, K., and D.M. Standley, 2013 MAFFT multiple sequence alignment software version 7: improvements in performance and usability. Molecular Phylogenetics and Evolution 30(4): 772–780.

Kim, D., L. Song, F.P. Breitweiser, and S.L. Salzberg, 2016 Centrifuge: rapid and sensitive classification of metagenomic sequences. Genome Research 26(12): 1721–1729.

Krehenwinkel, H., M. Wolf, J.Y. Lim, A.J. Rominger, W.B. Simison, et al., 2017 Estimating and mitigating amplification bias in qualitative and quantitative arthropod metabarcoding. Scientific Reports 7: 17668. https://doi.org/10.1038/s41598-017-17333-x

Kolmogorov, M., J. Yuan, Y. Lin, and P.A. Pevzner, 2019 Assembly of long, error-prone reads using repeat graphs. Nature Biotechnology 37: 540–546. https://doi.org/10.1038/s41587-019-0072-8

Legrand, B., J. Lesobre, J. Colombet, D. Latour, and M. Sabart, 2016 Molecular tools to detect anatoxin-a genes in aquatic ecosystems: Toward a new nested PCR-based method. Harmful Algae 58: 16–22.

Lepère, C., A. Wilmotte, and B. Meyer, 2000 Molecular diversity of *Microcystis* strains (Cyanophyceae, Chroococcales) based on 16S rDNA sequences. Systematics and Geography of Plants 70(2): 275–283

Li, H, 2018 Minimap2: pairwise alignment for nucleotide sequences. Bioinformatics 34(18): 3094–3100. https://doi.org/10.1093/bioinformatics/bty191

Martí, J.M., and C.P. Garay, 2019 Not just BLAST nt: WGS database joins the party. bioRxiv 653592. https://doi.org/10.1101/653592 (Preprint posted June 4, 2019)

Menegon, M., C. Cantaloni, A. Rodriguez-Prieto, C. Centomo, A. Abdelfattah, et al., 2017 On site DNA barcoding by nanopore sequencing. PLOS ONE 12(10): e0184741. https://doi.org/10.1371/journal.pone.0184741

Nguyen, L.-T., H.A. Schmidt, A. von Haeseler, and B.Q. Minh, 2015 IQ-TREE: A fast and effective stochastic algorithm for estimating maximum likelihood phylogenies. Molecular Biology and Evolution 32: 268–274. https://doi.org/10.1093/molbev/msu300

Pfeiffer, F., C. Gröber, M. Blank, K. Händler, M. Beyer, et al., 2018 Systematic evaluation of error rates and causes in short samples in next-generation sequencing. Scientific Reports 8: 10950. https://doi.org/10.1038/s41598-018-29325-6

Pomerantz, A., N. Peñafiel, A. Arteaga, L. Bustamante, F. Pichardo, et al., 2018 Real-time DNA barcoding in a rainforest using nanopore sequencing: opportunities for rapid biodiversity assessments and local capacity building. Gigascience 7: 1–14.

Quast, C., E. Pruesse, P. Yilmaz, J. Gerken, T. Schweer, et al., 2013 The SILVA ribosomal RNA gene database project: improved data processing and web-based tools. Nucleic Acids Research 41: D590–D596.

Rang, F.J., W.P. Kloosterman, and J. de Ridder, 2018 From squiggle to basepair: computational approaches for improving nanopore sequencing read accuracy. Genome Biology 19: 90. https://doi.org/10.1186/s13059-018-1462-9

Schafran, G.C, 2005 Reservoir Management Techniques to Improve Raw Water Quality. Algal Metabolytes Workshop, December 9, 2005, Sarasota, FL.

Seppey M., M. Manni, and E.M. Zdobnov, 2019 BUSCO: Assessing Genome Assembly and Annotation Completeness in Gene Prediction. Methods in Molecular Biology vol 1962, edited by M. Kollmar. Humana, New York, NY. doi.org/10.1007/978-1-4939-9173-0_14.

Sherwood, A.R., and G.G. Presting, 2007 Universal primers amplify a 23S rDNA plastid marker in eukaryotic algae and cyanobacteria. Journal of Phycology 43(3): 605–608. https://doi.org/10.1111/j.1529-8817.2007.00341.x

Stern, R.F., R.A. Andersen, I. Jameson, F.C. Küpper, M.A. Coffroth, et al., 2012 Evaluating the ribosomal internal transcribed spacer (ITS) as a candidate dinoflagellate barcode marker. PLOS ONE 7(8): e42780. https://doi.org/10.1371/journal.pone.0042780

Suurnäkki, S., G.V. Gomez-Saez, A. Rantala-Ylinen, J. Jokela, D.P. Fewer, et al., 2015 Identification of geosmin and 2-methylisoborneol in cyanobacteria and molecular detection methods for the producers of these compounds. Water Research 68: 56–66.

Tsao, H.-W., A. Michinaka, H.-K. Yen, S. Giglio, P. Hobson, et al., 2014 Monitoring of geosmin producing *Anabaena circinalis* using quantitative PCR. Water Research 49:416–425. http://dx.doi.org/10.1016/j.watres.2013.10.028.

Vaser, R., I. Sović, N. Nagarajan, and M. Šikić, 2017 Fast and accurate de novo genome assembly from long uncorrected reads. Genome Research 27: 737–746. https://doi.org/10.1101/gr.214270.116

Wacklin P., L. Hoffmann, and J. Komárek, 2009 Nomenclatural validation of the genetically revised cyanobacterial genus *Dolichospermum* (Ralfs ex Bornet et Flahault) comb. nova. Fottea 9(1): 59–64.

Wick, R.R., L.M. Judd, and K.E. Holt, 2019 Performance of neural network basecalling tools for Oxford Nanopore sequencing. Genome Biology 20: 129. https://doi.org/10.1186/s13059-019-1727-y

Wick R.R., M.B. Schultz, J. Zobel, and K.E. Holt, 2015. Bandage: interactive visualisation of *de novo* genome assemblies. Bioinformatics 31(20): 3350–3352.

Wongsurawat T., M. Nakagawa, O. Atiq, H.N. Coleman, P. Jenjaroenpun, et al., 2019 An assessment of Oxford Nanopore sequencing for human gut metagenome profiling: A pilot study of head and neck cancer patients. Journal of Microbiological Methods 166: 105739. https://doi.org/10.1016/j.mimet.2019.105739

Wood, D.E., J. Lu, and B. Langmead, 2019 Improved metagenomic analysis with Kraken2. Genome Biology 20: 257.

Zhang, Y., X. Lin, T. Li, H. Li, L. Lin, et al., 2020 High throughput sequencing of 18S rRNA and its gene to characterize a *Prorocentrum shikokuense* (Dinophyceae) bloom. Harmful Algae 94: 101809.

Zhu, W., A. Lomsadze, and M. Borodovsky, 2010 *Ab initio* gene identification in metagenomic sequences. Nucleic Acids Research 38(12): e132. https://doi.org/10.1093/nar/gkq275

